# Single-cell proteomics of pre-implantation mouse embryos uncovers distinct asymmetry of certain proteins among early blastomeres

**DOI:** 10.1101/2024.09.28.614398

**Authors:** Yuan Yuan, Mo Hu, Yinghui Zheng, Yutong Zhang, Yuxuan Pang, Xiaoliang Sunney Xie

## Abstract

During pre-implantation mouse embryonic development, blastomeres undergo division and differentiation, reaching a distinctive level of heterogeneity, hence the completion of the first cell fate determination. However, when the initial asymmetry emerges and how this heterogeneity amplifies, particularly at the protein level, remain elusive. Here, by mass spectrometry-based single-blastomere proteomics, we identified proteins exhibiting significant heterogeneity in abundance among mouse blastomeres as early as the 2-cell stage. Differential gene expression among blastomeres, as indicated by intra-embryo variation in RNA abundance detected through single-cell RNA sequencing, was insufficient to fully explain the corresponding disparities in protein abundance. Instead, the asymmetric distribution of protein molecules during cell division was observed, serving as another mechanism contributing to protein heterogeneity, independent of RNA expression.

## Introduction

Following the union of gametes, the fertilized egg initiates a process of differentiation and gives rise to distinct cell lineages, making the inception of life in mammals. Understanding when heterogeneity arises among the zygote’s progenies and how the asymmetry evolves constitute central inquiries in developmental biology. In many organisms, such as *C. elegans*, the asymmetric pre-patterning of the embryo is pronounced, resulting in explicit heterogeneity among blastomeres and distinct cell fate determinations. In contrast, this phenomenon is not applicable in the context of mammalian pre-implantation embryos. For example, mouse embryos initially appear to be uniform, displaying no overt morphological distinction among blastomeres until the fourth division. It is at this stage that some blastomeres adopt inner positions, while others remain on the embryo’s periphery(1, 2). In the subsequent one to two rounds of cell division, the first cell lineage specification segregates blastomeres into the inner cell mass (ICM), destined to form the new organism and the yolk sac, and the trophectoderm (TE), which will give rise to the placenta. A hypothesis has been made which suggests that in early-stage mammalian embryos, before the emergence of inside and outside cells, blastomeres are equivalent in terms of fate and potency(3–9).

On the contrary, recent research in mouse embryonic development suggests that the activity of specific cell fate regulators, and hence the potency of blastomeres, may exhibit asymmetry as early as the 4-cell stage (10–12). With the advent of single-cell omics technology, analyses of single-cell transcriptome(12–24), translatome(25, 26), epigenome(27–35) and 3D chromatin architecture(36, 37) have greatly deepened our understanding of pre-implantation mammalian embryonic development. Many studies support the perspective that blastomeres in 4-cell and even 2-cell stage mammalian embryos are indeed different from each other at the transcriptomic or epigenomic level(21, 22, 28, 33, 38).

On the flip side, proteins serve as executors in most biological processes, directly participating in developmental progression. Considering that protein abundance often cannot be accurately predicted by mRNA abundance(39, 40), the direct assessment of protein levels in early mammalian embryos stands as an imperative endeavor to advance our understanding of embryonic development. Previous investigations have offered valuable insights into the regulatory mechanisms governing embryogenesis through bulk proteomic profiling of pre-implantation mouse embryos(41–44). However, these bulk analyses yield averaged protein levels across embryo populations, impeding our ability to discern inter-blastomere differences.

The recent advances in mass spectrometry (MS)-based proteomics(45–52) have enabled the quantification of proteomic profiles at the single-blastomere level in several studies(52–56). However, these studies are either confined to specific stages of the pre-implantation developmental process(53, 55, 56) or do not prioritize the comparative analysis of protein abundance among blastomeres originating from the same embryo(52, 54). A comprehensive examination of the single-cell level protein expression landscape during pre-implantation mouse embryonic development, with a focus on when and how heterogeneity among blastomeres from the same embryo arises and expands, is still lacking.

Herein, using a label-free quantitative MS strategy, we analyzed the protein expression landscape of mouse embryos in-depth at the single-blastomere level, spanning from the zygote to the morula stage. Our results unveil proteins that exhibit conspicuous difference in abundance between blastomeres derived from the same two-cell and four-cell stage embryos. We also performed single-blastomere RNA-seq experiments and detected intra-embryo RNA asymmetry at various stages, consistent with previous studies. However, we found that variations in RNA abundance, indicative of differential gene expression among blastomeres within the same embryo, were insufficient to explain the corresponding heterogeneity in protein abundance, suggesting the involvement of additional mechanisms. We demonstrate that the asymmetric distribution of protein molecules into daughter blastomeres during cell division serves as an RNA-expression-independent mechanism underlying the generation and amplification of blastomere-to-blastomere heterogeneity.

## Results

### Overview of the experiment

Utilizing the MS-based label-free single-cell proteomic pipeline we recently developed(51), we embarked on the examination of proteomic profiles of individual blastomeres across five developmental stages, ranging from the zygote to the morula stage (Table. S1). Embryos at various developmental stages were dissociated into single blastomeres, with each blastomere retaining its embryo assignment (Fig. 1A). Blastomeres were then digested and analyzed one-by-one using an Orbitrap Eclipse mass spectrometer. In total, we analyzed 291 blastomeres and identified 2,978 proteins, of which 283 blastomeres and 2,703 proteins passed the quality control. Notably, we consistently detected around 1,000 proteins in the majority of single-cell samples, despite the progressive halving of cell volumes during early blastomere divisions(57) (Fig. S1A). Uniform manifold approximation and projection (UMAP) analysis successfully separated blastomeres from different developmental stages into clusters (Fig. 1B).

**Fig. 1.**
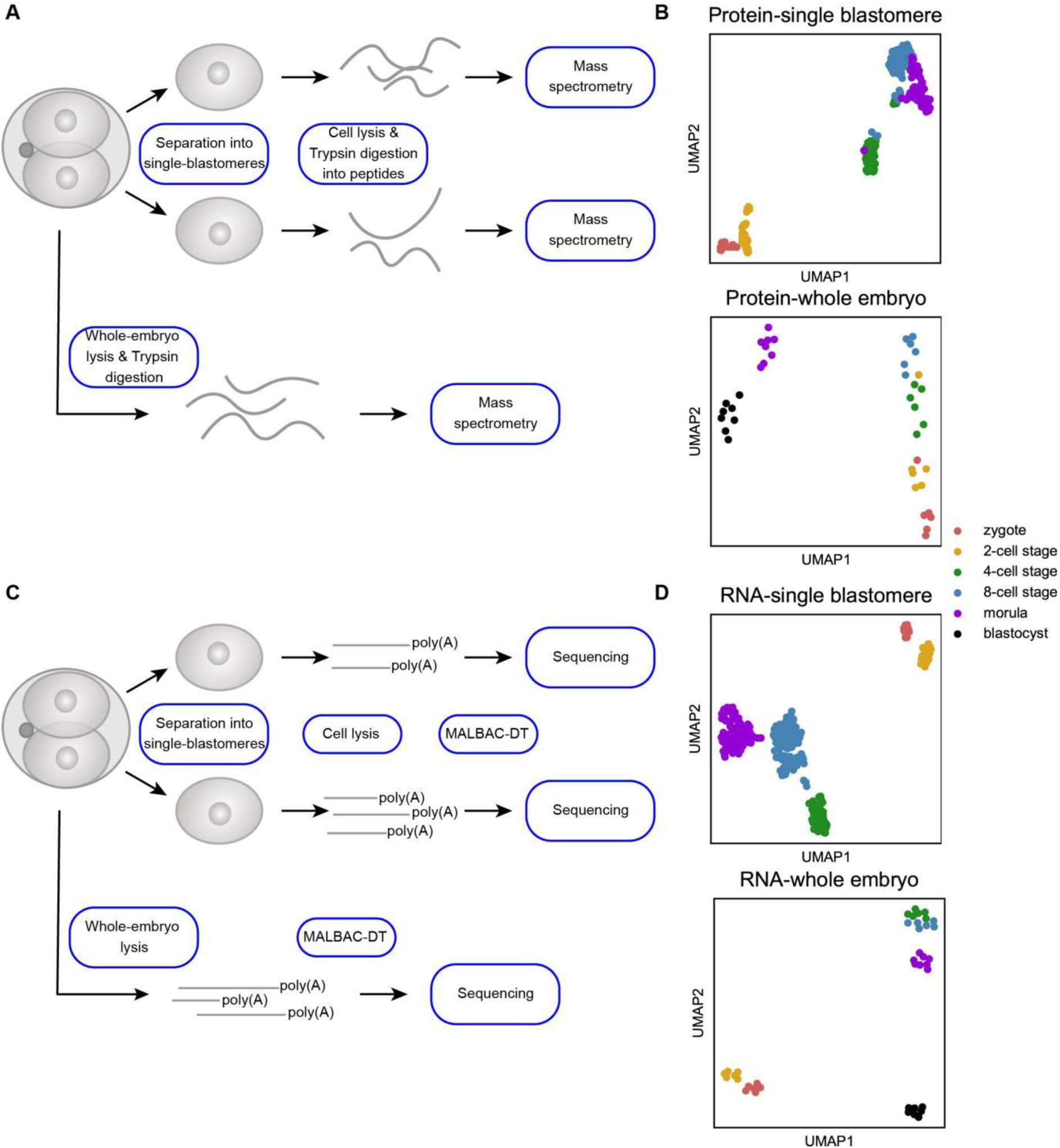
Overview of the experiment. (A) Mouse embryos at various developmental stages were separated into single blastomeres while preserving their respective embryo identities. Blastomeres were then subjected to digestion, and protein abundance was measured by mass spectrometry-based label-free proteomics. Another set of samples were analyzed one-by-one as whole embryos. (B) Uniform manifold approximation and projection (UMAP) analysis of single blastomeres (n=283) and whole embryos (n=40) at different stages based on the proteomic data. Each dot represents an individual blastomere or embryo. (C) Similar to (A), but with transcriptomic analysis conducted via multiple annealing and looping based amplification cycles for digital transcriptomics (MALBAC-DT). (D) UMAP analysis of single blastomeres (n=258) and whole embryos (n=39) at different stages based on the transcriptomic data.

For single-blastomere transcriptomic analysis, we dissociated embryos using the same approach and employed multiple annealing and looping based amplification cycles for digital transcriptomics (MALBAC-DT)—a single-cell RNA-seq method developed by our group, which provides high RNA quantification accuracy(58) (Fig. 1C). A total of 258 blastomeres were analyzed, resulting in the identification of around 10000 genes in each sample (Fig. S2A).

To delineate the gene expression profile of whole embryos across various pre-implantation embryonic stages, complementing single-blastomere datasets, we conducted a single-embryo proteomic profiling of 40 embryos spanning from the zygote to the blastocyst stage using the same MS-based analytical pipeline and detected more than 2,000 proteins in most samples (Fig. S1B). We also performed a transcriptomic assessment of other 39 embryos using MALBAC-DT. UMAP analysis of the transcriptomic and proteomic datasets of whole embryos also distinguished embryos at varying developmental stages from each other (Fig. 1B and D), demonstrating the high quality of our data.

### Whole-embryo transcriptomic and proteomic profiling exhibits major events in early embryogenesis

Through the integration of whole-embryo transcriptomic and proteomic datasets, we sought to elucidate the gene expression landscape during pre-implantation mouse embryonic development. To this end, we examined the dynamic changes in gene expression occurring between each successive developmental stage. We focused on significantly up- or down-regulated genes among overlapped genes between the transcriptomic and the proteomic analysis (Fig. 2A). As a consequence of zygotic genome activation (ZGA) (57), hundreds of genes were up-regulated at the RNA level during the developmental progression from zygotes to 2-cell stage embryos. Concurrently, a substantial decline in RNA abundance of another group of genes was observed, likely attributable to the degradation of maternal RNAs. Intriguingly, our analysis revealed that despite the pronounced decrease in RNA levels, only a minor subset of corresponding proteins showed a significant reduction in their abundance. This result underscores that maternal RNA degradation does not guarantee a corresponding decline in protein abundance, aligning with previous studies(42, 44).

**Fig. 2.**
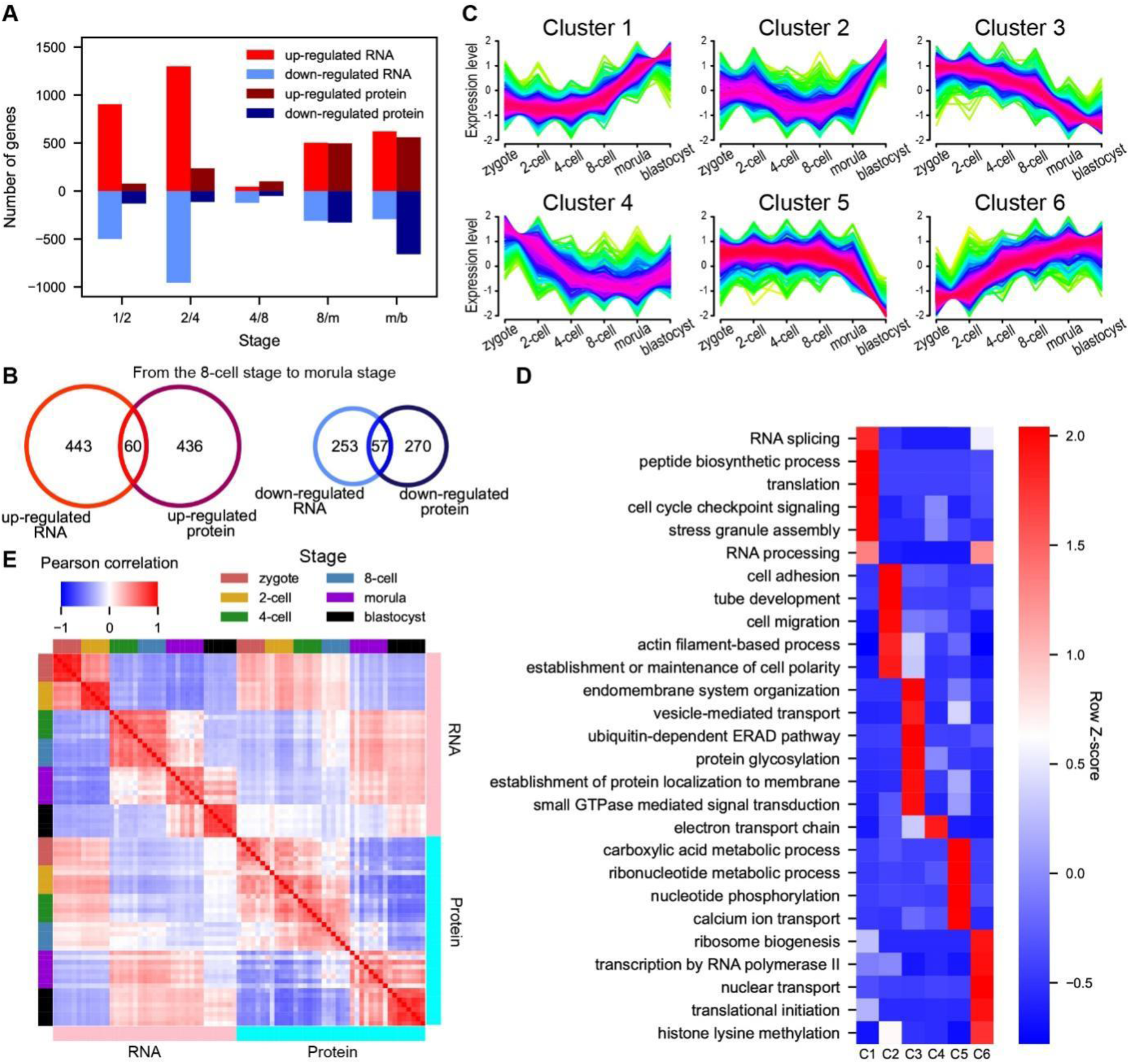
Comprehensive gene expression landscape of whole mouse embryos. (A) Bar plot illustrating the numbers of significantly up- or down-regulated RNAs and proteins between two consecutive stages (m=morula, b=blastocyst). A gene is considered significantly altered if the fold change between the averaged RNA UMI count or protein abundance of two consecutive stages is greater than 2 or smaller than 0.5, and the p-value is smaller than 0.05, Welch’s t-test. Only genes simultaneously detected in whole-embryo transcriptomic and proteomic analyses are shown. (B) Venn plot depicting the overlap of up- or down-regulated genes between RNA and protein abundance measurements during the transition from 8-cell stage embryos to morulae. (C and D) Identification of six distinct temporal patterns of protein expression by fuzzy c-means clustering (C), and Gene Ontology biological process (GOBP) analysis of proteins from each cluster (D). For each GOBP term, the averaged enrichment p-value across six clusters is subtracted from the p-value of each cluster, then divided by the standard deviation of p-values for that term to obtain the Z-score. (E) Pearson correlation coefficient of scaled whole-embryo RNA and protein abundance. RNA and protein levels are separately scaled by subtracting the averaged abundance of each gene across all samples in each dataset and dividing by the standard deviation of the abundance of that gene.

What’s more, a drastic increase in expression levels of many genes commenced from the 8-cell stage of embryonic development. While similar amounts of genes were up-regulated at RNA and protein levels during the transition from the 8-cell stage embryo to the morula, the genes identified as up-regulated from these two datasets hardly overlapped with each other (Fig. 2B, left). A similar trend was also discerned among genes that experienced down-regulation during this developmental progression, demonstrating that changes at the RNA level could not accurately predict alterations in protein abundance.

In order to systematically investigate the temporal dynamics of proteins throughout pre-implantation mouse development, we applied fuzzy c-means algorithm(59) and grouped all detected proteins into six clusters, each showing different temporal trends (Fig. 2C). For instance, components of the subcortical maternal complex (SCMC)(60), including MATER, FILIA, FLOPED, TLE6, PADI6 and ZBED3, were all classified into cluster 5. This is consistent with previous findings which indicate that SCMC components endure through early developmental stages and undergo degradation in the blastocyst stage(61). Epigenomic modifiers were also detected. TET3, which is responsible for mediating global demethylation soon after fertilization(62), displayed a consistently down-regulated pattern and was assigned to cluster 3. *De novo* methyltransferases DNMT3A/B were allocated to cluster 4 and 2, respectively. Noteworthy was the remarkable up-regulation of DNMT3B in blastocysts, possibly getting ready for the re-establishment of the DNA methylation landscape after implantation(63). Temporal trends of transcription factors (TFs) also featured prominently. Specifically, the trophectodermal specifier CDX2 exhibited a heightened expression level at the blastocyst stage (cluster 2). Similarly, the totipotent lineage specifier OCT4 displayed up-regulation in later stages (cluster 1). SALL4 and TEAD4, acting upstream of OCT4 and CDX2 respectively, mirrored the trend observed in cluster 1. Gene Ontology biological process (GOBP) analysis of each cluster (Fig. 2D) revealed that up-regulated proteins exhibited an enrichment in RNA and protein synthesis processes, aligning with previous results(42). Up-regulated proteins were also found to be responsible for the formation of cell adhesion and polarity. In contrast, down-regulated proteins were associated with metabolic processes, endomembrane system organization and protein localization.

To gain deeper insights into the extent of dependence of protein levels on RNA abundance in pre-implantation mouse embryos, we analyzed the Pearson correlation coefficient for scaled whole-embryo RNA and protein abundance among embryos at each developmental stage (Fig. 2E). While embryos at the same stage showed good uniformity in both transcriptomic and proteomic profiles, the RNA-protein correlation within each stage could be rather poor. This observation further demonstrates that variances of RNA abundance do not necessarily translate into corresponding alterations in protein abundance. Modulations of translation rate or differences between RNA and protein half-lives can effectively mitigate the impact of changes at the transcript level. Interestingly, we also found that the transcriptomes from 4- and 8-cell stage embryos exhibited superior correlations with the proteomes of morula and blastocyst stage embryos than with those of embryos at the same stage. This finding aligns with prior researches that it may take time for newly synthesized RNAs to be translated, thus changes of protein abundance are lagged behind(40, 44). The correlation between the two datasets, based on genes rather than samples, exhibited a similar trend (Fig. S3).

### Single-cell proteomic analysis reveals inter-blastomere heterogeneity at the protein level in early pre-implantation mouse embryos

When the initial asymmetry becomes discernible in mouse embryos, especially at the protein level, remains a longstanding question. Based on our single-blastomere proteomic dataset spanning five pivotal stages before implantation, we probed blastomere-to-blastomere heterogeneity, if any, using all proteins within our detection scope. Applying a previously published equation(64, 65), we ascertained that the overall protein expression bias showed a continual increase as embryonic development progresses (Fig. 3A), which is not surprising as blastomeres gradually lose their totipotency and embark on different cell fate trajectories.

**Fig. 3.**
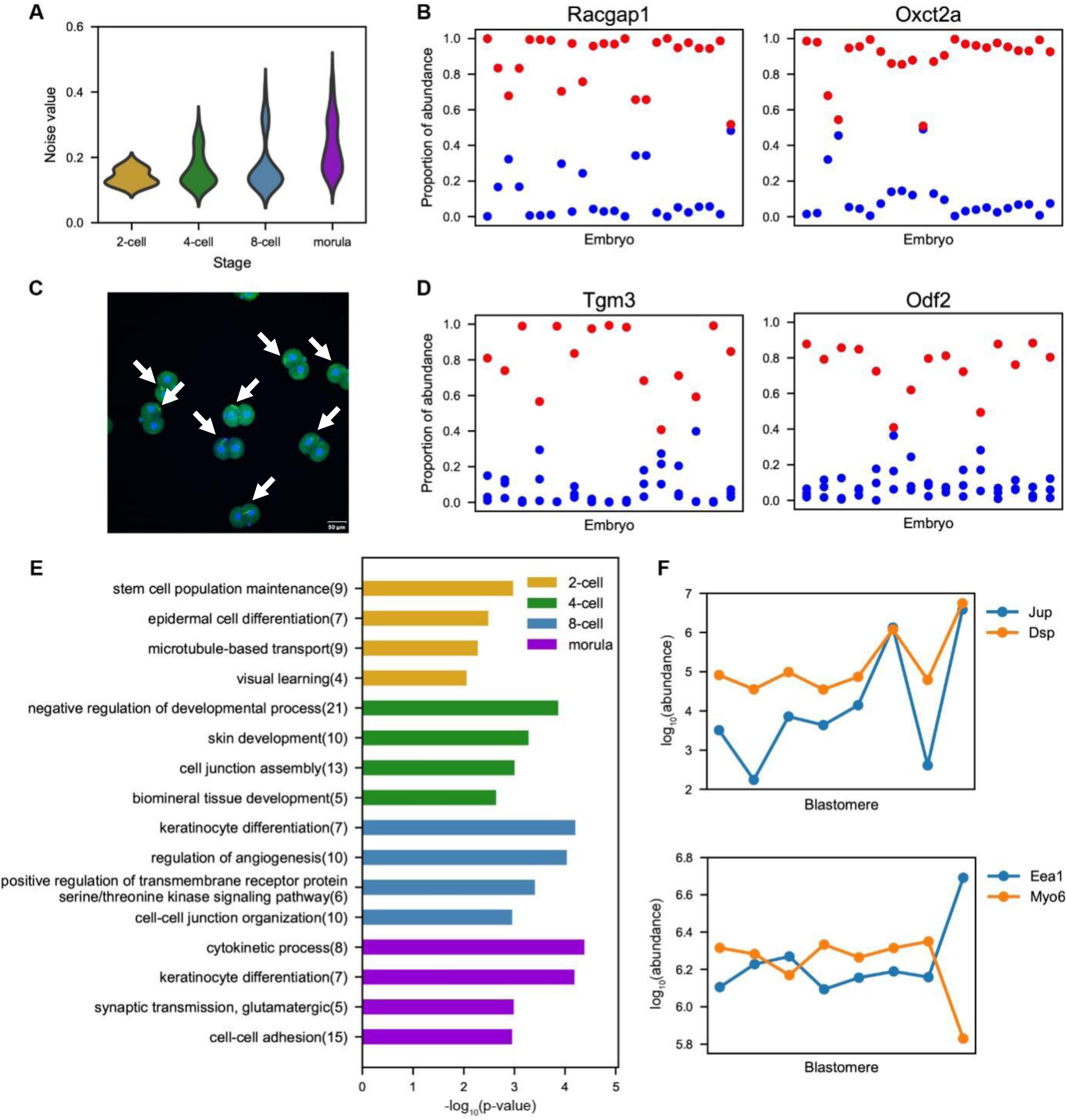
Single-blastomere proteomic analysis reveals blastomere-to-blastomere heterogeneity in mouse embryos. (A) Violin plot showing noise values between blastomeres within the same embryo at different stages. (B and D) Examples of proteins exhibiting a bimodal distribution pattern among blastomeres in 2-cell (B) and 4-cell (D) stage embryos. Each dot represents a blastomere, and dots in the same column originate from the same embryo. (C) A confocal image of an immunofluorescence assay of OXCT2A (Alexa Fluor 488) in 2-cell stage mouse embryos. DAPI fluorescence indicates cell nuclei. (E) Gene Ontology biological process (GOBP) analysis of differentially abundant proteins (DAPs) at each developmental stage. DAPs are proteins for which the averaged intra-embryo coefficient of variation (CV) of abundance across all embryos at the corresponding stage ranks in the top 200 among proteins detected in at least half of the blastomeres at that stage. Top enriched terms are listed, and the number of proteins involved in each term is shown in parentheses. (F) Example of protein pairs whose levels are correlated (top) or anti-correlated (bottom) among blastomeres in an 8-cell stage embryo.

Subsequently, our investigation delved into whether there were individual proteins exhibiting consistent disparities among blastomeres derived from the same early-stage embryo. We utilized the method published previously(21) and partitioned the differences in protein abundance between blastomeres into two distinct components: intra-embryo and between-embryo variation. We discovered that certain proteins displayed striking heterogeneity between blastomeres at the 2-cell stage, with their cell-to-cell variation mainly arising from intra-embryo variation. For instance, the abundance of RACGAP1 and OXCT2A within blastomeres of 2-cell stage embryos exhibited a distinct bimodal distribution pattern (false discovery rate<0.001, Benjamini-Hochberg procedure) (Fig. 3B). There were also proteins whose abundance within one of the blastomeres in 4-cell stage embryos markedly surpassed that in the other three blastomeres, such as TGM3 and ODF2 (false discovery rate<0.01, Benjamini-Hochberg procedure) (Fig. 3D). Our comprehensive analysis revealed a set of 484 proteins exhibiting larger intra-embryo variation than between-embryo variation within 2-cell stage embryos, reflecting the nontrivial asymmetry of protein abundance.

We further categorized proteins as differentially abundant proteins (DAPs) at each developmental stage. These proteins exhibited a significant intra-embryo coefficient of variation (CV) (top 200) among proteins detected in at least half of the blastomeres at their respective stages. Our analysis identified 9 proteins that showed overlapping presence in DAPs of all four developmental stages: OXCT2A, RACGAP1, EIF4ENIF1, TGM1, TGM3, BLMH, HSPB1, HTT and LMAN2L. We posit that these proteins play crucial roles in the initiation and amplification of inter-blastomere bias, thereby contributing to the process of cell fate determination.

We then performed the GOBP analysis of DAPs at each stage, yielding insights into their functional attributes (Fig. 3E). For instance, we observed that DAPs became significantly enriched in processes related to cell junctions, especially proteins actively involved in cell-cell adhesion, both structurally (PLEC, JUP, PKP1, DSP, CTNNA1 etc.) and functionally (CDC42, ROCK2 etc.), starting from the 4-cell stage. This observation suggests a potential association between cell-cell interactions and the process of blastomere differentiation. Our analysis revealed that at the 2-cell and the 4-cell stages, the top-ranking GO terms enriched among DAPs were “stem cell population maintenance” and “negative regulation of developmental process”, respectively. In particular, the inclusion of ICM markers like SALL4 and CARM1 in DAPs at the 2-cell and the 4-cell stages, respectively, suggests the potential existence of different pluripotency among blastomeres as early as the 2-cell stage. Proteins involved in “cytokinetic process” were enriched at the morula stage. This result might be attributed to the divergence in cell division patterns during the transition from 8-cell to morula stage embryos, encompassing the symmetric division, generating two outer cells, and the asymmetric division, giving rise to one inner cell and one outer cell.

Beyond the identification of proteins exhibiting distinct abundance among blastomeres within the same embryo, we supposed that protein pairs sharing structural or functional relationships would demonstrate synchronous alterations in abundance. After removing the between-embryo variation(21), we calculated Pearson correlation coefficients for protein pairs across blastomeres at each stage. Indeed, our investigation revealed multiple protein pairs characterized by high positive correlations. For instance, we found that JUP and DSP, both components of desmosome(66), displayed a strong correlation within 8-cell stage embryos (Fig. 3F, r=0.62). At the same time, many protein pairs were found to exhibit strikingly negative correlations, such as EEA1 and MYO6 (Fig. 3F, r=-0.74). A prior study revealed that EEA1, a marker of endosome, was more abundant in MYO6-deficient epithelial cells. This phenomenon was attributed to the diminished efficiency of endosome-lysosome fusion-a process promoted by MYO6(67). This possibly explains the observed negative correlation between these two proteins. Interestingly, EEA1-positive early endosomes have been reported to be asymmetrically distributed in *C. elegans* embryos depending on actomyosin contractility and embryonic polarity(68). The asymmetric spatial distribution of endosomes has been hypothesized to play a role in cell fate determination within *C. elegans* embryos. Whether a similar pattern of endosome distribution occur in pre-implantation mouse embryos and whether such distribution exerts an influence in embryonic development require further study.

### Comprehensive single-blastomere transcriptomic and proteomic analysis uncovers discrepancies between RNA and protein asymmetry in pre-implantation mouse embryos

We then explored how protein asymmetry is established and progresses. One straightforward mechanism suggests that differential gene expression across blastomeres results in heterogeneity in RNA abundance, subsequently leading to protein asymmetry. To test this hypothesis, we conducted single-blastomere RNA-seq experiments to investigate whether a similar heterogeneity exists at the RNA level within mouse embryos. To reduce the possible influence of technical noise in single-cell RNA-seq experiments(22), we focused on a subgroup of genes with high RNA abundance (top 50%) for further analysis. We identified RNAs showing clear expression bias within embryos (Fig. S5A and B). Notably, certain cell fate specifiers, such as Sox21 and Nanog, exhibited differential abundance across blastomeres, which is in agreement with previous findings(12). We also defined differentially abundant RNAs at each developmental stage. Biological processes like “transmembrane receptor protein serine/threonine kinase signaling pathway” were enriched among these RNAs (Fig. S5C). Additionally, we found that RNA abundance generally varies more significantly between blastomeres compared with protein levels (Fig. 4A). This may result from shaper fluctuations in RNA abundance due to their shorter half-lives and lower median molecule counts relative to proteins.

**Fig. 4.**
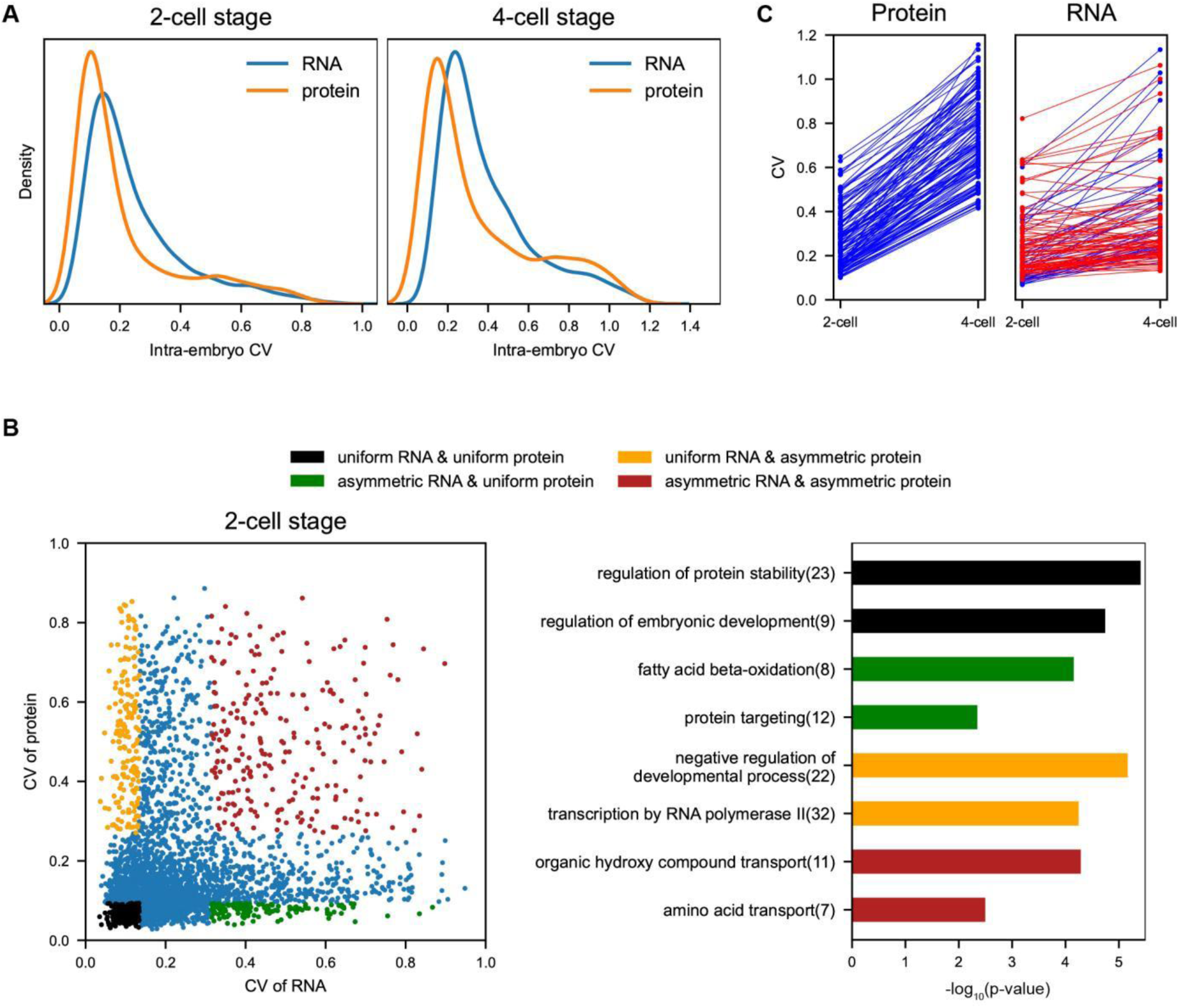
Comparison of RNA and protein asymmetry within pre-implantation mouse embryos. (A) Kernel density estimate (KDE) plot of the intra-embryo CV for RNAs and proteins at the 2-cell and the 4-cell stages. Overall, RNAs exhibit greater variation in abundance between blastomeres compared with proteins. (B) Scatter plot showing the extremely low correlation (Pearson r=0.09) between the intra-embryo CV of RNA and the intra-embryo CV of protein at the 2-cell stage (left), and GOBP analysis of genes showing either similar or distinct abundance uniformity at RNA and protein levels within embryos (right). (C) Changes in the intra-embryo CV of proteins with mean abundance per blastomere increased from the 2-cell to the 4-cell stage, accompanied by a significant increase in CV (CV at the 4-cell stage is at least 0.3 greater than that at the 2-cell stage) (left), and CV changes of corresponding RNAs (right). Blue lines (n=38) indicate a significant increase in the intra-embryo CV of RNA, while red lines (n=83) represent cases where the CV of RNA did not change significantly (Benjamini-Hochberg adjusted p-value < 0.05, Welch’ s t-test) or even declined, exhibiting a divergent trend from that of the protein.

To better elucidate the relationship between RNA and protein heterogeneity, we analyzed an additional 93 blastomeres from the 2-cell and the 4-cell stages using the timsTOF SCP mass spectrometer, harvesting higher proteome coverage. Through the integrative analysis of single-blastomere transcriptomic and proteomic datasets, we aimed to explore the correlation between RNA and protein asymmetry-specifically, whether RNA abundance heterogeneity results in corresponding protein abundance bias, in accordance with the RNA-dependent mechanism. Intriguingly, we observed a near-zero correlation (Pearson r=0.09) between the intra-embryo CV of RNA and that of protein at both the 2-cell (Fig. 4B) and 4-cell stage (Fig. S7), highlighting the discordance between RNA and protein asymmetry. Notably, we identified genes exhibiting distinct differences in abundance uniformity between RNA and protein levels within embryos. Some differentially abundant proteins at the 2-cell stage, including EZH2 and PTPN2, which were enriched in “negative regulation of developmental process”, displayed uniform RNA abundance among blastomeres. These differences in protein abundance would have been overlooked if we focused solely on RNA heterogeneity, which could not directly link to functional biological processes. This underscores the importance of our single-blastomere proteomic analysis in highlighting the asymmetry of protein distribution.

We also evaluated whether differential gene expression serves as the primary mechanism driving protein heterogeneity amplification during the transition from the 2-cell to the 4-cell stage. We selected a subgroup of proteins whose mean abundance per blastomere increased during the transition, accompanied by a significant increase in intra-embryo CV at the protein level. According to the mechanism, these genes would be differentially expressed across blastomeres in 4-cell stage embryos, exhibiting greater RNA heterogeneity compared with the 2-cell stage. This RNA heterogeneity would then lead to increased protein abundance and asymmetry, consistent with the selection criteria for these proteins. However, we found that more than half of the corresponding RNAs did not show a significant change in intra-embryo CV during this transition (Fig. 4C), with some even showing a decrease in CV. These results demonstrate that differential gene expression among blastomeres is insufficient to account for the observed disparities in protein abundance, indicating that additional mechanisms are crucial in establishing protein asymmetry.

### Integration of whole-embryo and single-blastomere datasets reveals asymmetric distribution of proteins as an important mechanism underpinning the establishment of intra-embryo heterogeneity

Asymmetric distribution of specific transcripts and proteins have been found to play a central role in embryonic development and cell fate determination(69–71). We recognize that the asymmetric distribution of protein molecules during cell division may also lead to blastomere-to-blastomere bias independent of RNA expression. For instance, the RNA abundance of Gstk1, a glutathione transferase, remains consistently low throughout the pre-implantation embryonic development, while its whole-embryo protein abundance is maintained at a relatively constant level. As a result, the expression of Gstk1 is largely inactive, and differential gene expression has been excluded as a contributing factor. However, the CV of the protein within embryos at each stage continuously increases (Fig. 5B). This increase in abundance heterogeneity could be attributed to the uneven distribution of protein molecules during cell division, which represents a key mechanism in addition to the RNA-dependent differential gene expression.

**Fig. 5.**
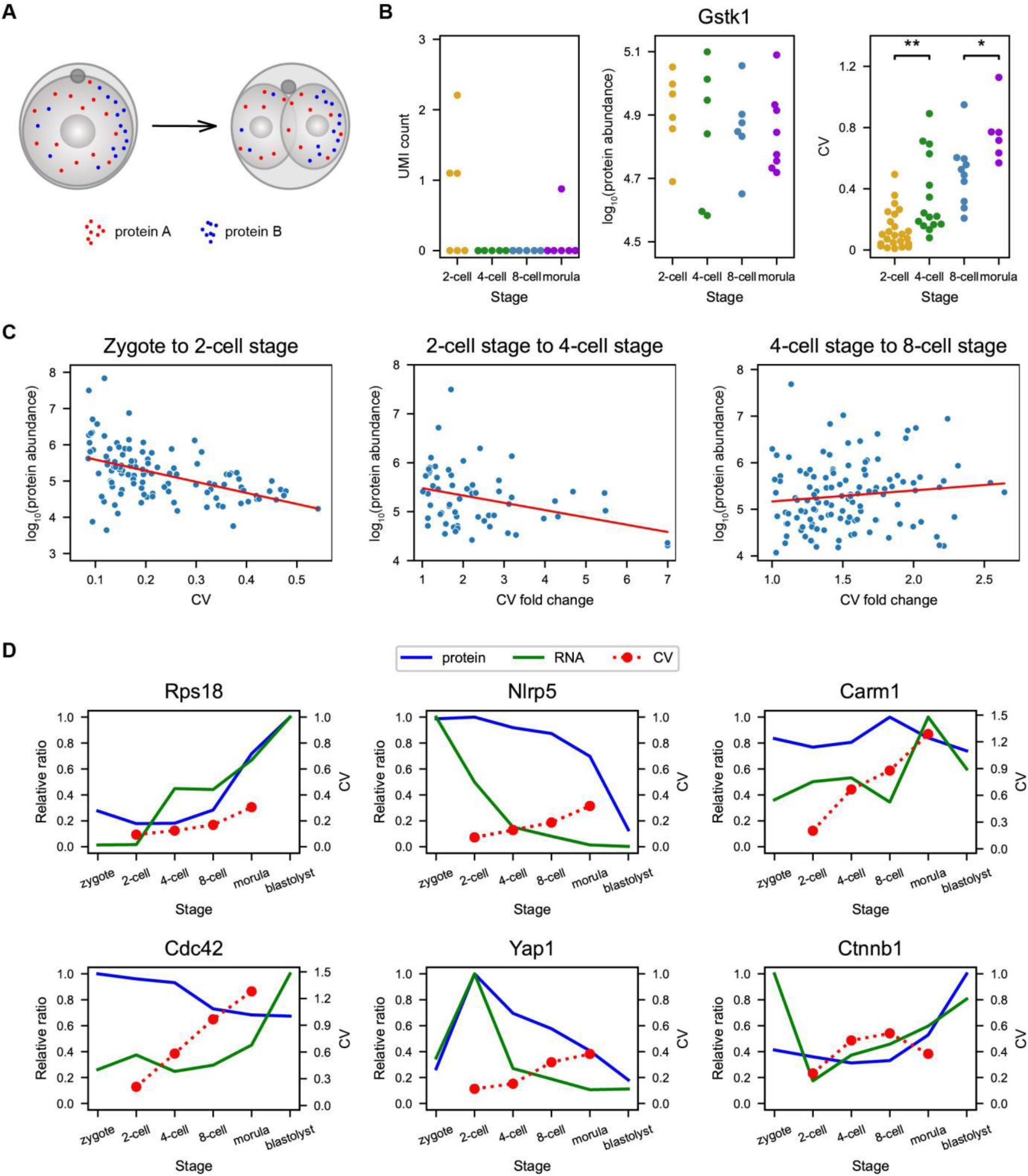
Asymmetric distribution of protein molecules during cell division is a pivotal mechanism in forming heterogeneity among blastomeres. (A) Schematic diagram illustrating that protein molecules can be either symmetrically (protein A) or asymmetrically (protein B) distributed into two daughter cells during cell division. (B) An example of a protein for which intra-embryo heterogeneity is primarily influenced by asymmetric distribution. Each dot represents corresponding data generated from an embryo. *p-value<0.05, **p-value<0.01, Tukey’s HSD test. (C) Plots showing the correlation between the averaged protein abundance and the averaged intra-embryo CV of proteins in 2-cell stage embryos (left), between the averaged protein abundance and the averaged CV fold change during the transition from the 2-cell to the 4-cell stage (middle), and from the 4-cell to the 8-cell stage (right). Each dot represents a protein. For each selected protein, its averaged normalized UMI counts of the corresponding RNA in whole embryos at both consecutive stages are less than 4, its change in averaged whole-embryo protein abundance between the two stages is less than 20%, and the protein is detected in at least half of blastomeres at both stages. (D) Examples of the expression landscape of genes throughout pre-implantation embryonic development. For the averaged RNA UMI count and protein abundance in whole embryos at each stage of a specific gene, the largest value among stages is scaled to 1, and values of other stages are converted relative to the largest value, shown according to the left longitudinal axis of each subplot (proteins in blue lines and RNAs in green lines). Averaged absolute intra-embryo CV value across embryos at each stage is shown according to the right longitudinal axis (red dotted lines).

Motivated by this finding, we investigated genes with mostly silent expression and stable protein abundance, attributing changes in variation between consecutive stages primarily to the distribution of proteins among daughter cells. Typically, if protein molecules are randomly partitioned into two daughter cells, the probability of asymmetric partitioning increases with decreasing protein abundance. This phenomenon was indeed observed during the first embryonic division (Fig. 5C, left), as the intra-embryo CV of protein abundance in 2-cell stage embryos showed a negative correlation with their abundance (Pearson r=-0.49). A similar trend was also discovered during the transition from the 2-cell to the 4-cell stage (Fig. 5C, middle, Pearson r=-0.34). Notably, near half of the proteins selected during this transition showed a significant increase in CV (p-value<0.05, Benjamini-Hochberg procedure), indicating that, in some cases, the mechanism of asymmetric distribution of protein molecules alone is sufficient to establish and amplify protein asymmetry.

Furthermore, we identified a shift in this pattern during the transition from 4-cell to 8-cell stage embryos. Here, the CV fold change exhibited a slightly positive correlation with the overall protein abundance in whole embryos (Fig. 5C, right, Pearson r=0.12), indicating that deterministic processes predominate over random distribution in this developmental phase. This is interesting as it aligns with a hypothesis that stochastic events play an important role in creating early heterogeneity, while deterministic regulation becomes more prominent as development progresses(22). Our investigation reveals that the asymmetric distribution of protein molecules during cell division is an RNA-independent mechanism contributing to protein heterogeneity, in addition to differential gene expression among blastomeres.

Through the integration of whole-embryo transcriptomic, proteomic and single-blastomere proteomic data, we have achieved a comprehensive depiction of the expression landscape of each detected gene throughout embryonic developmental process (Fig. 5D). Different proteins show distinct expression patterns depending on the categories of their coding genes and their roles in pre-implantation mouse embryonic development. For instance, the housekeeping gene Rps18 was up-regulated at both RNA and protein levels, especially in later stages, supporting the needs for embryonic growth, whereas the inter-blastomere variation of the protein RPS18 was constantly low. The maternal protein NLRP5 also displayed low variation between blastomeres, with a rapid decline in abundance at the blastocyst stage because the gene should not be expressed shortly after fertilization and the corresponding protein is degraded in later stages.

The expression profiles of cell fate specifiers exhibited distinct patterns compared with housekeeping genes. For instance, the RNA and protein abundance of ICM marker Carm1 remained relatively stable throughout development. However, the protein exhibited increasing asymmetry between blastomeres, demonstrating its crucial role in biasing cells towards distinct fates. CDC42 is a Rho GTPase known for actively promoting polarization. It has been reported that depletion of CDC42 diminishes Cdx2 expression and thus blastomeres tend to commit to the ICM cell fate(72). Our analysis revealed an elevated intra-embryo bias without substantial fluctuation in the whole-embryo protein level, akin to the behavior of CARM1. YAP1 acts as a central regulator of cell fate through the Hippo signaling pathway. When the Hippo signaling pathway is activated, YAP1 is phosphorylated and remains in the cytoplasm. When the pathway is deactivated, unphosphorylated YAP1 enters the nucleus, binds with its partner TEAD4 and promotes the expression of many TE marker genes such as Cdx2(73). We found that the protein variation between blastomeres remained low, aligning with its role in determining cell fate through cellular localization rather than abundance bias. There were also many proteins that exhibited a reduction in inter-blastomere variation at certain developmental stages. For example, beta-catenin, responsible for cell-cell adhesion and participating in the Wnt signaling pathway, displayed up-regulation in abundance from the 8-cell stage onward, accompanied by a decreased bias among blastomeres. The elucidation of the reason behind the declined CV at the morula stage requires further investigation. Expression profiles of more genes are provided in Fig. S8.

## Discussion

The emergence and progression of heterogeneity among blastomeres in the mouse embryo, particularly at the protein level, represent pivotal inquiries in the field of developmental biology. Gaining deeper insights into these questions holds the potential to significantly advance reproductive and stem cell research. Our single-blastomere proteomic data, obtained from a MS-based experimental pipeline, revealed that certain proteins exhibited a bimodal distribution in embryos as early as the 2-cell and the 4-cell stages. While it may be surprising to observe pronounced heterogeneity in protein abundance at such early stages, previous studies have claimed that progenies of 2-cell and 4-cell stage blastomeres contribute nonuniformly to ICM and TE cells in the blastocyst (74–80). However, these results offer limited depth in elucidating molecular mechanisms governing mammalian embryonic development. Other studies delved more deeply into the molecular-level examination of blastomere heterogeneity at the 4-cell stage. The heterogeneity has been ascribed to the different abundance of CARM1, a histone-arginine methyltransferase, and consequently to the levels of H3R26 methylation among blastomeres(10). Higher H3R26me levels were posited to enhance the binding affinity of transcription factors such as Oct4 and Sox2 to DNA, thereby promoting the transcription of pluripotent genes(12).

Among DAPs identified at the 2-cell and the 4-cell stages, there are noteworthy other candidates for further investigation. For instance, RACGAP1, recognized as a component of the centralspindlin complex involved in cell division, has also been found to participate in the differentiation of hematopoietic cell(81). This prompts us to consider the potential role of RACGAP1 in blastomere differentiation. Signaling pathways reliant on cell adhesion may also contribute to the establishment of heterogeneity among blastomeres. For example, the Wnt signaling pathway, in which some DAPs such as JUP participate, may play a role in this process.

Differential gene expression among blastomeres is an essential and straightforward mechanism driving protein asymmetry. However, our findings underscore that it is not the sole mechanism at play. The asymmetric distribution of protein molecules into daughter cells during cell division serves as an RNA-independent mechanism that establishes and amplifies heterogeneity among blastomeres, and in some cases, may even be the dominant source of the heterogeneity. What’s more, while the notion that more abundant proteins exhibit an even distribution aligns with our intuitive expectations in early stages, a reversal of this trend was observed during the transition from the 4-cell to the 8-cell stage. The observed asymmetric distribution of proteins during this phase appears more to be a regulated process rather than a stochastic one.

How exactly the deterministic allocation of proteins into one of the daughter cells happens remains an open question. We propose two possible mechanisms. Firstly, we consider the spatially asymmetric distribution of protein molecules within the mother cell, building on the notion that even eggs and zygotes have been proposed to exhibit spatial inhomogeneity(1, 2). Secondly, we hypothesize that active transport of proteins by cytoskeletal filaments play a role in directing proteins into a specific daughter cell during cell division, as we discovered that certain actin-binding proteins, such as PFN2, exhibit both high abundance and a significant increase in CV during the transition from 4-cell to 8-cell embryos (Fig. 5C).

In summary, our single-blastomere proteomic analysis has unveiled the emergence of protein heterogeneity among mouse embryo blastomeres at the protein level as early as the 2-cell stage. Differential gene expression among blastomeres is not the sole factor contributing to disparities in protein abundance. Asymmetric distribution of protein molecules during cell division can also emerge as the dominant source of protein asymmetry. Our study provides a valuable resource for understanding of pre-implantation mouse embryonic development at the molecular level. Further investigation on mechanistic intricacies of how protein heterogeneity manifests in mouse embryos, along with elucidating biological functions they confer, will deepen our comprehension of the developmental process.

## Acknowledgments

This project is financially supported by Beijing Advanced Innovation Center for Genomics at Peking University and the Beijing Municipal Commission of Science and Technology (Z201100005320015). We thank Dr. Wenping Ma for his guidance on performing MALBAC-DT experiments. We thank Dr. Fangnong Lai for helpful suggestions on the experimental procedure. We also thank Dr. Wenyang Dong, Dr. Honggui Wu and Fanchong Jian for insightful discussions on the project, and staff of High-throughput Sequencing Center at Peking University and Mass Spectrometry Core at Changping Laboratory for their help in sample sequencing and analysis.

## Author contributions

Y. Y., M. H., and X. S. X. designed the experiments; Y. Y., M. H., Y. Zheng and Y. Zhang collected the experimental data; Y. Y., M. H., and Y. P. analyzed the data; and Y. Y., M. H., and X. S. X. wrote the manuscript with the help of all authors.

---

**Fig. S1.**
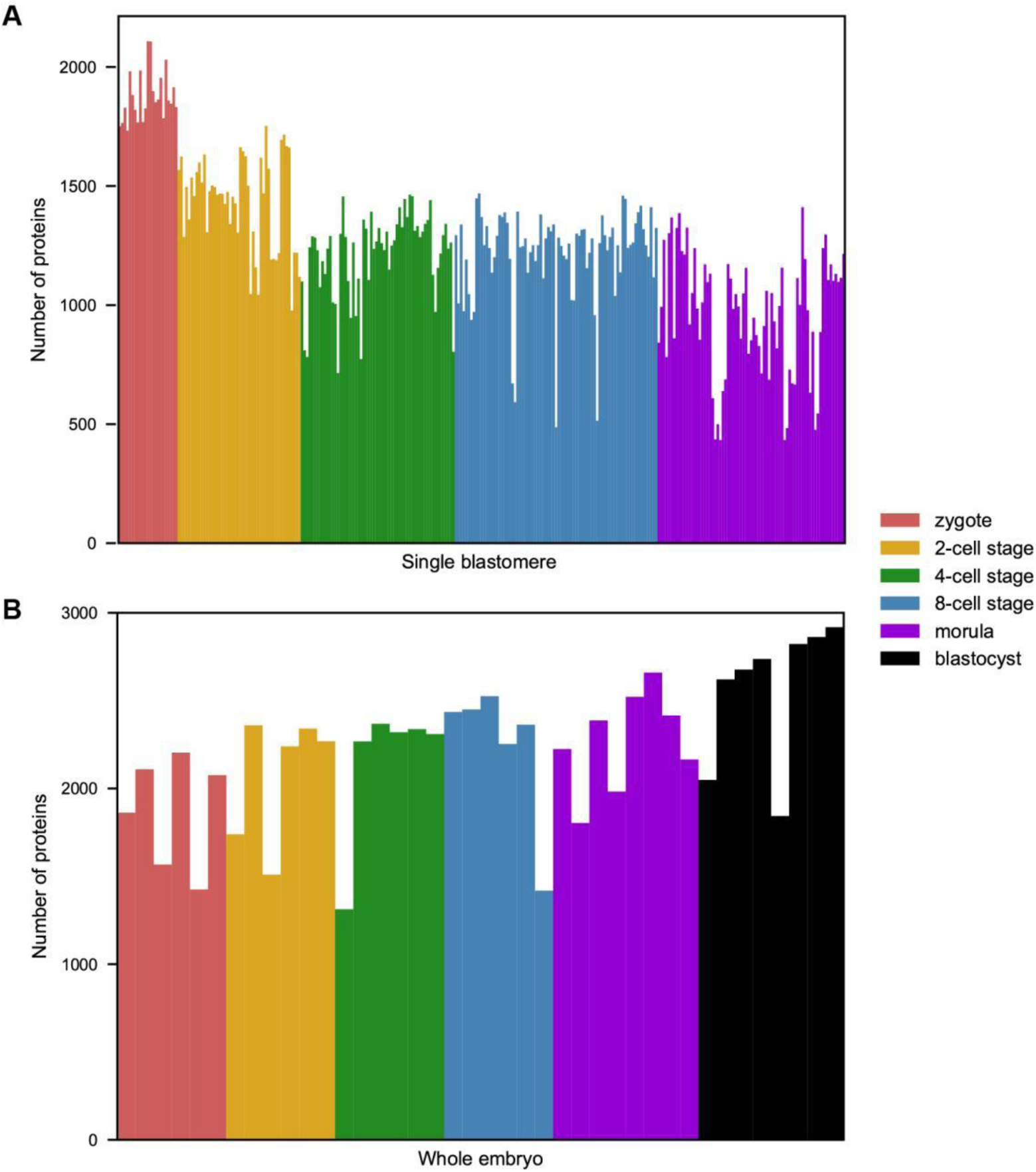
Numbers of proteins detected in whole embryos and blastomeres at various developmental stages. Over 2000 proteins were detected in most whole embryos. More than 1000 proteins were identified in the majority of single blastomeres as well.

**Fig. S2.**
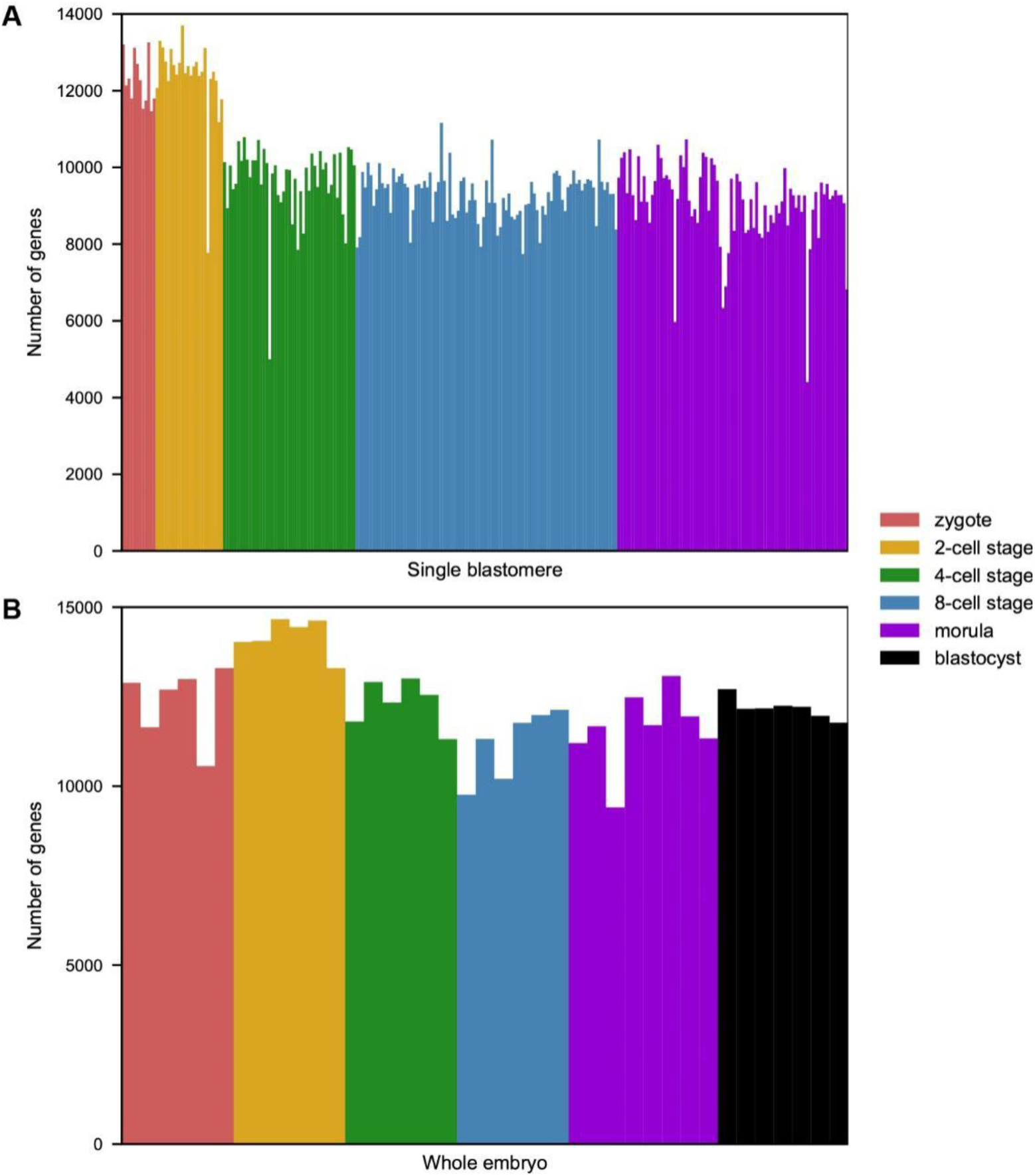
Identification numbers of genes in whole embryos and blastomeres at various developmental stages. Over 10000 genes were identified in most samples.

**Fig. S3.**
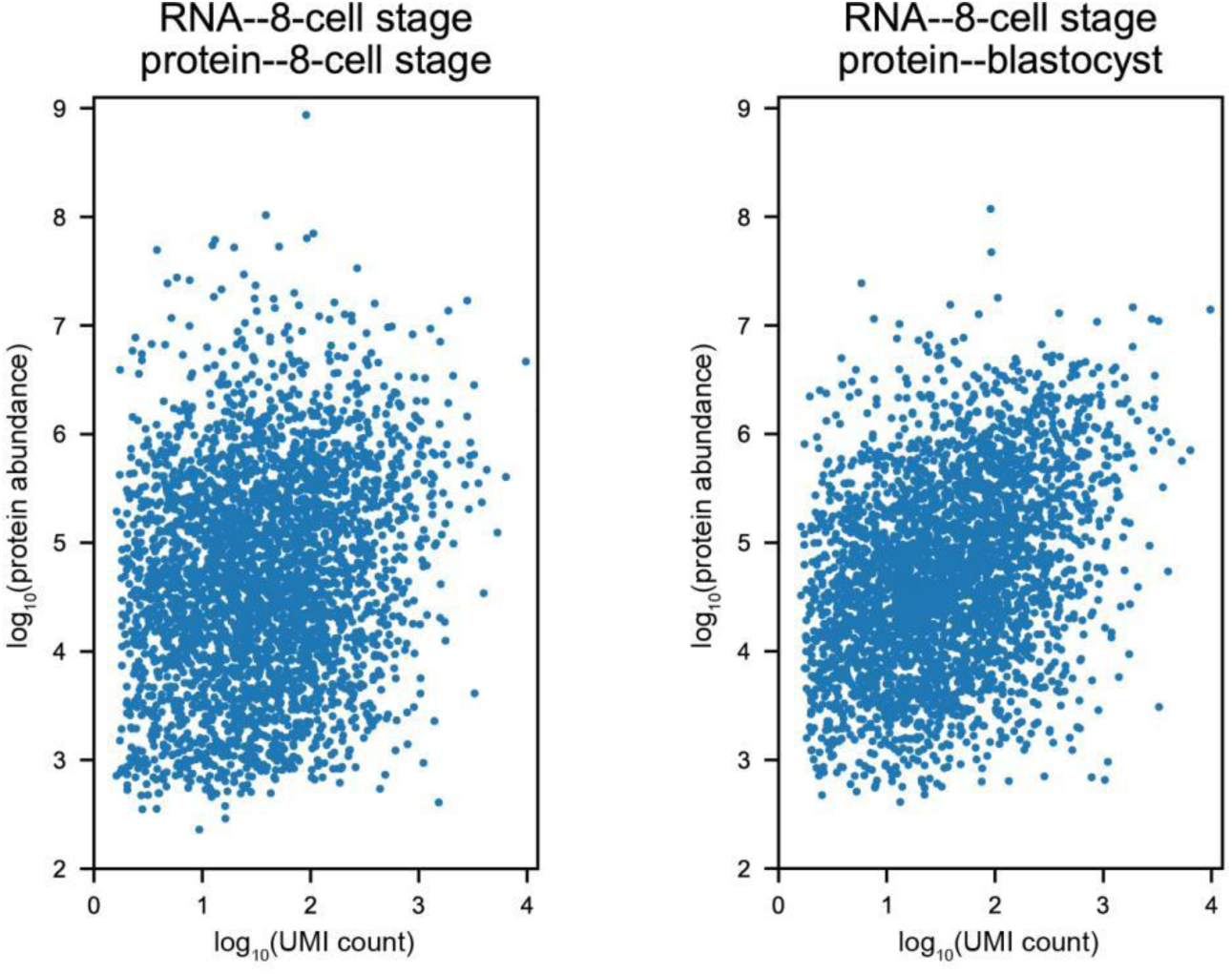
Scatter plot illustrating correlations between RNA UMI counts of 8-cell stage embryos and protein abundance in 8-cell stage embryos (left, Pearson r=0.20) or in blastocysts (right, Pearson r=0.35). Each dot represents a gene. RNA UMI counts of embryos at earlier stages were found to be better correlated with protein abundance of later-stage embryos, demonstrating the “phase-shift” between RNA and protein abundance changes.

**Fig. S4.**
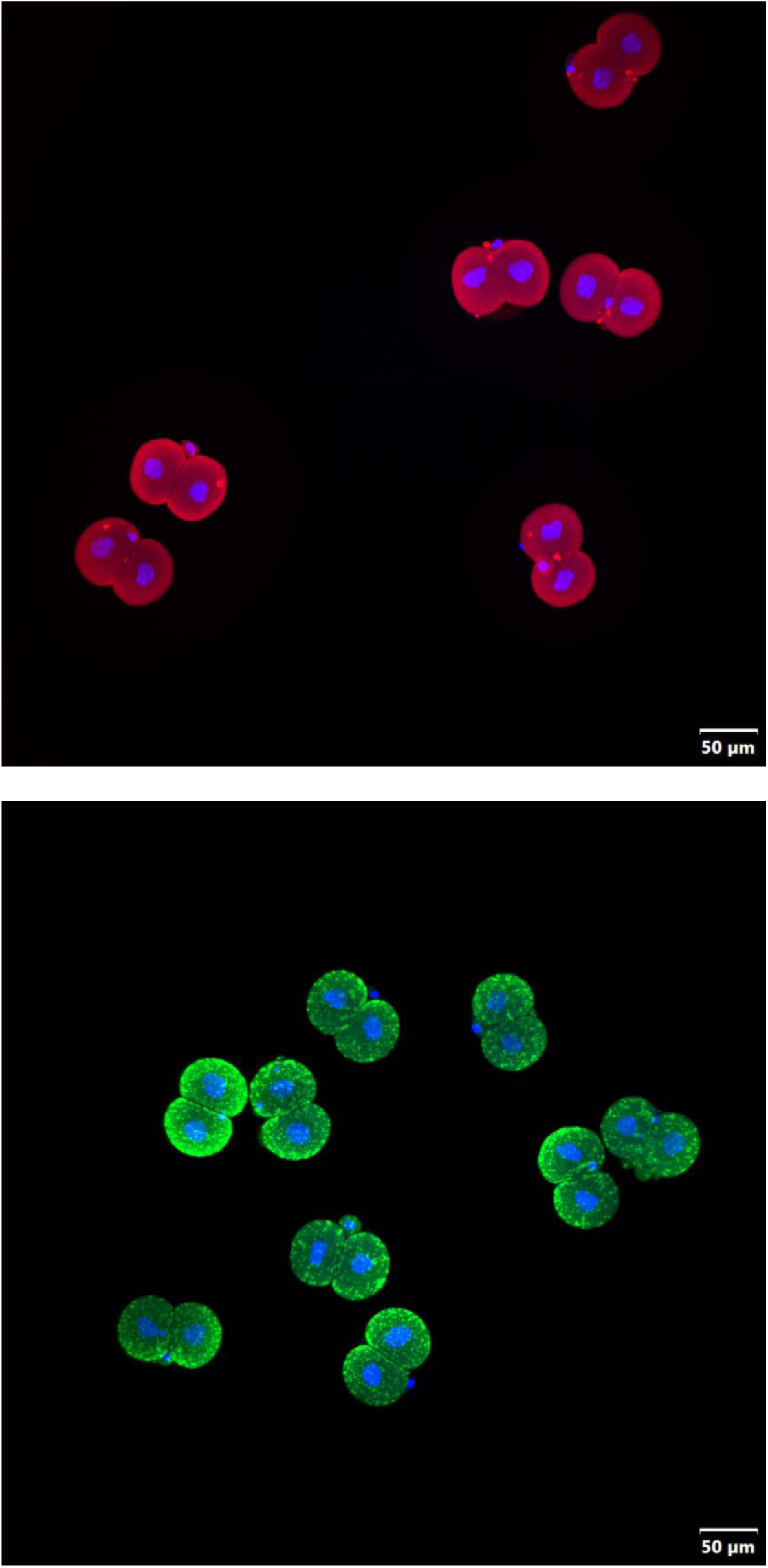
Confocal images of an immunofluorescence assay of RACGAP1 (top, Alexa Fluor 568) and TGM3 (bottom, Alexa Fluor 488). DAPI fluorescence indicates cell nuclei. Differentially abundant proteins were validated by distinct fluorescence intensities between blastomeres.

**Fig. S5.**
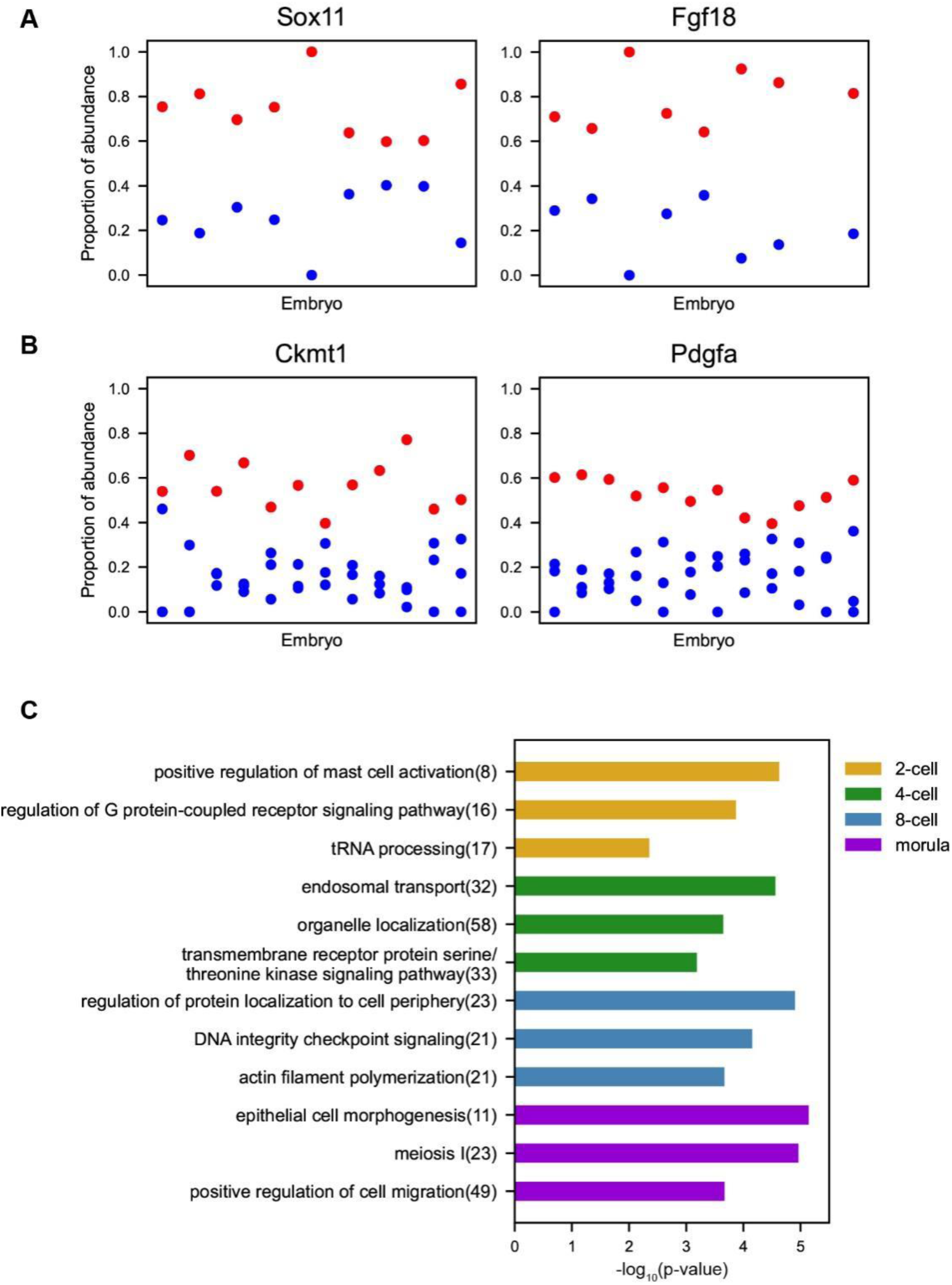
(A-B) Examples of RNAs exhibiting a bimodal distribution pattern among blastomeres in 2-cell (A) and 4-cell (B) stage embryos. Each dot represents a blastomere, and dots in the same column originate from the same embryo. (C) GOBP analysis of differentially abundant RNAs at each developmental stage. These RNAs show strong expression bias (top 1000 by intra-embryo CV among those in the top 50% of expression levels). Top enriched terms are listed, with the number of proteins involved in each term indicated in parentheses.

**Fig. S6.**
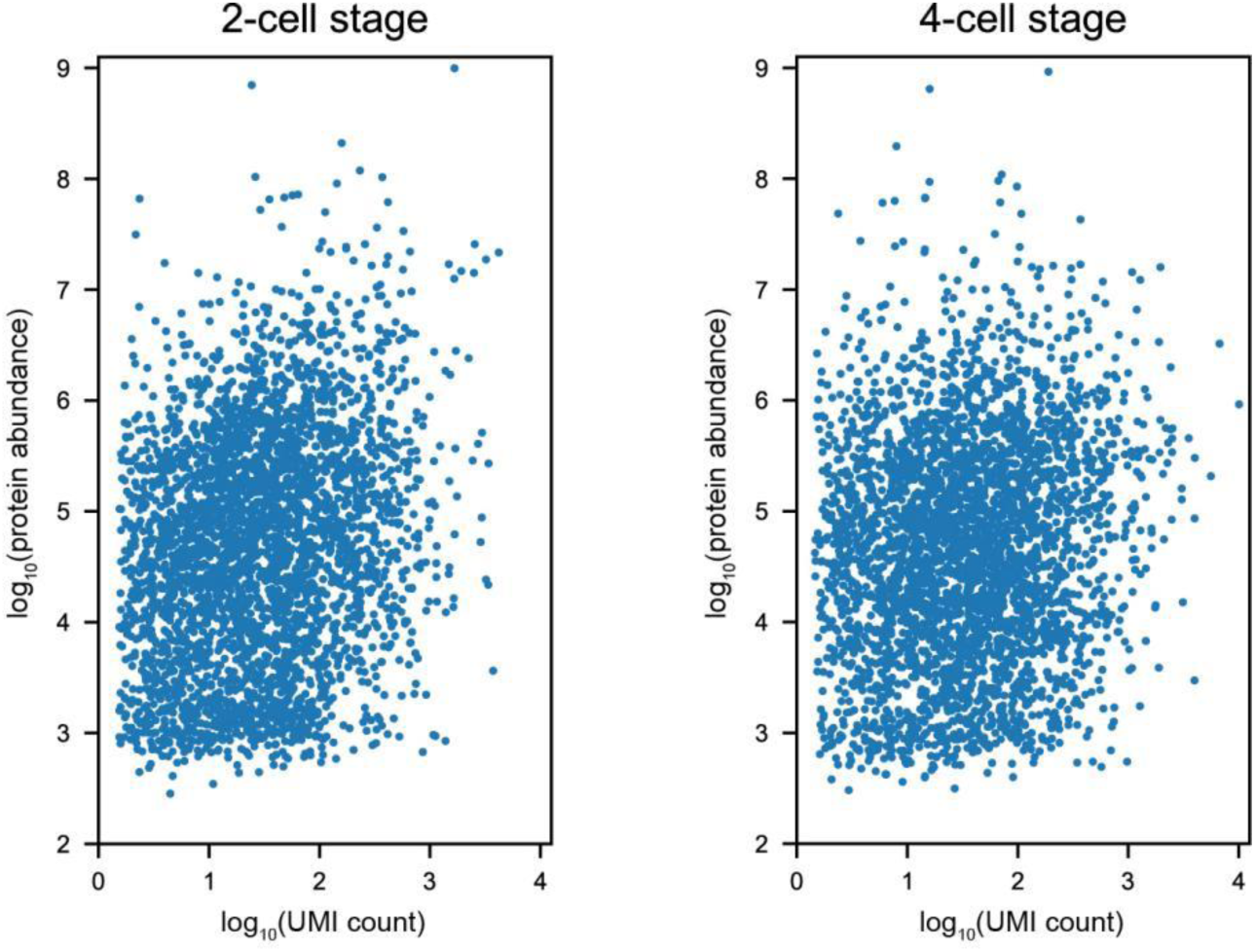
Scatter plot showing the correlation between RNA UMI counts and protein abundance in 2-cell stage embryos (left, Pearson r=0.21) and 4-cell stage embryos (right, Pearson r=0.13). Each dot represents a gene.

**Fig. S7.**
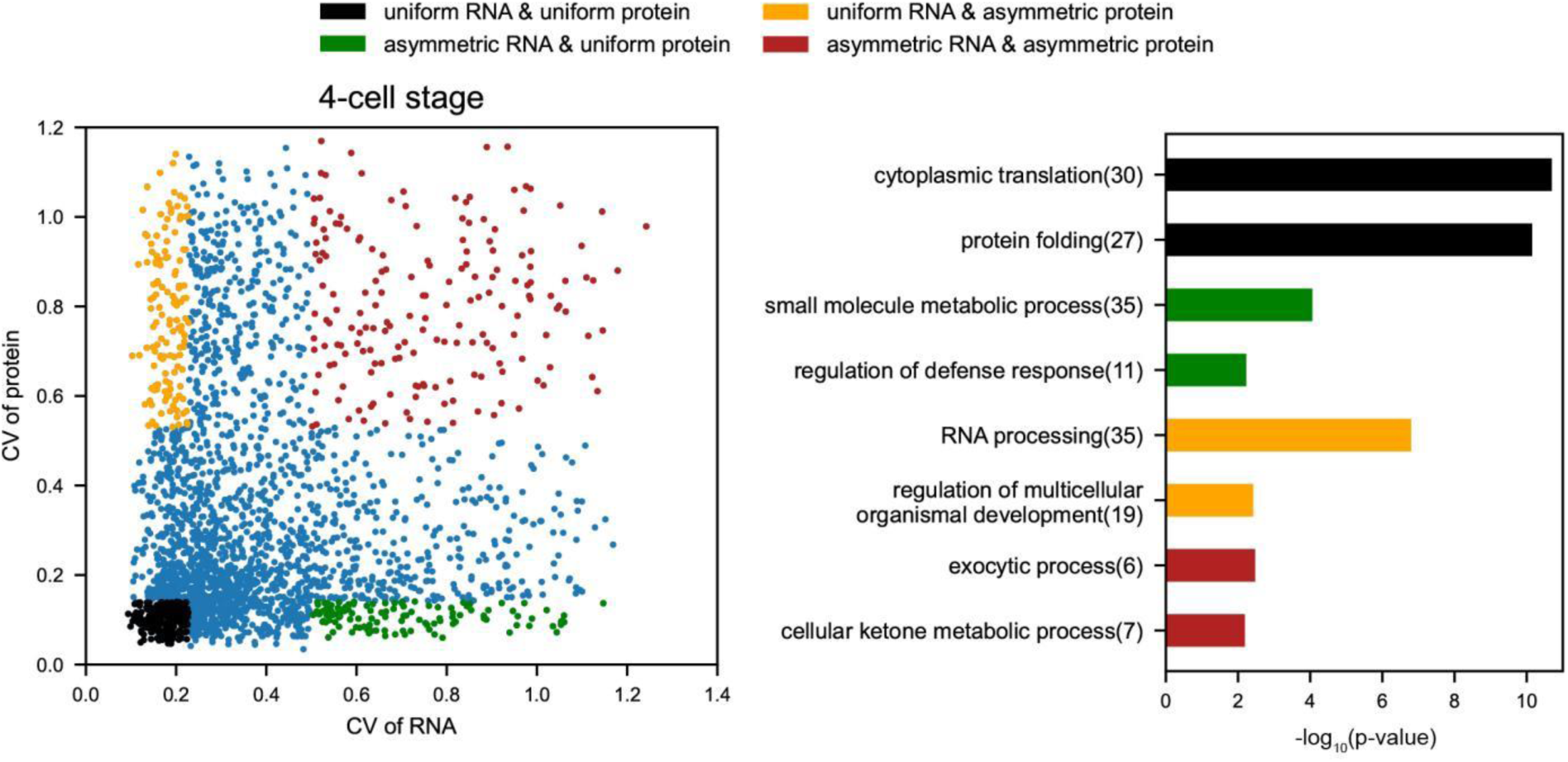
Scatter plot showing the correlation (Pearson r=0.09) between the intra-embryo CV of RNA and the intra-embryo CV of protein within 4-cell stage embryos (left), and GOBP analysis of genes showing either similar or distinct abundance uniformity at RNA and protein levels within embryos (right).

**Fig. S8.**
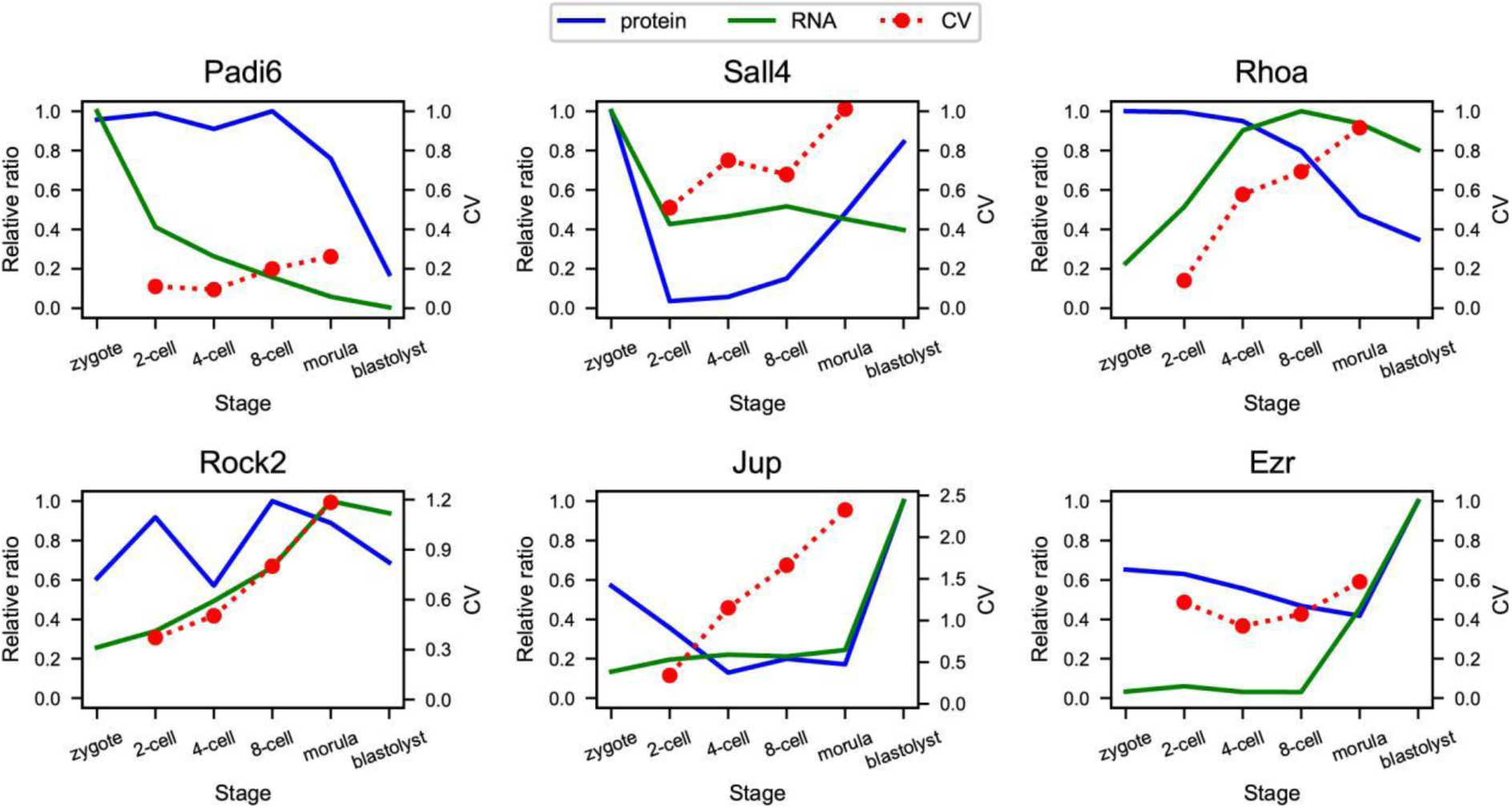
More examples of the expression landscape of genes throughout pre-implantation embryonic development.

**Table. S1.** Number of samples at each developmental stage.

| Stage | Zygote | 2-cell | 4-cell | 8-cell | Morula | Blastocyst |
| --- | --- | --- | --- | --- | --- | --- |
| Single-blastomere proteomics | 23 (23 embryos) | 48 (24 embryos) | 60 (15 embryos) | 79 (10 embryos) | 73 (6 embryos) |  |
| Whole-embryo proteomics | 6 | 6 | 6 | 6 | 8 | 8 |
| Single-blastomere transcriptomics | 12(12 embryos) | 24(12 embryos) | 47(12 embryos) | 93(12 embryos) | 82(6 embryos) |  |
| Whole-embryo transcriptomics | 6 | 6 | 6 | 6 | 8 | 7 |

